# Mosquitoes (Diptera:Culicidae) in an indigenous settlement of the Magdalena State, Colombia

**DOI:** 10.1101/2020.04.23.056101

**Authors:** Gabriel Parra-Henao, Erick Perdomo, Deibys Carrasquilla, Emy Torres, Celenny Perez, Jose Luis Molina

## Abstract

Studies directed to investigate ecological parameters of mosquitoes populations allows to establish the risk of vector borne diseases transmission and to bring recommendations for health authorities about prevention, surveillance and control. We report some ecological aspects of mosquitoes populations in a region of Colombia. Quarterly sampling was done. For mosquito catching CDC and Shannon traps were used. Diversity and abundance indexes estimation was done. A total of 1071 mosquitoes belonging to four genera and 10 species were collected. The most abundant species were *Cx quinquefasciatus* (22.6%), *Ae. aegypti* (20.08%), *Ae. scapularis* (12.3%), *Ae. angustivittatus* (9%) and *An. albimanus* (8.0%). The finding of *Cx pedroi, Ae. scapularis*,, *Cx. nigripalpus, Cx. quinquefasciatus, Cx. declarator, Ma. titillans*, and *Ps. ferox* previously reported as arbovirus vectors warns about the possibility of transmission in the zone and the finding of *An. albimanus*, also warns about the risk for malaria transmission in the area.

## Introduction

In Colombia, because of the expansion of farming and environmental modifications, the indigenous populations have been displaced to small areas called “resguardos” where new interactions with mosquito’s populations have emerged, leading to new epidemiological patterns and new scenarios of transmission of vector-borne diseases. There is growing evidence that demographic movements coupled with environmental alterations may favors the proliferation of vector-borne diseases, the main ones being malaria, lymphatic filariasis, encephalitis, and onchocerciasis (Nozais, 2003)

The indigenous population in Magdalena State are divided into four ethnic groups (Kogi, Arzario, Arhuaco, and Ette Enaka). The Ette Enaka group live in a settlement called Issa Oristuna that is surrounded by a matrix of modified landscape used mainly for cattle production.

In the Issa Oristuna settlement, no inventory of mosquitoes was done, but in states near this area (with a distance no less than 200 km) there are some description of mosquitoes (Rivas F et al, 1997; Ferro et al, 2008), These studies reported the presence of *Culex* (*Melanoconion*) *pedroi, Cx. (Mel) ocossa, Cx quinquefasciatus, Phosorophora confinis, Mansonia indubitans* and *Ae. taeniorhynchus*.

The knowledge of the geographic distribution of mosquito species in areas of indigenous settlements is important to planning and targeting epidemiological surveillance and vector control activities to protect the health of this vulnerable ethnic groups, for this reason, the aim of the present study is to report the first inventory of mosquitoes with epidemiological importance in the area of Issa Oristuna indigenous settlement.

## Methods

The study was performed from February to September 2018 in the village of Issa Oristuna, San Angel Municipality located in the Colombian North West (Caribbean region), in the geographic coordinates N 10.01555 and W −73. 305694’, annual precipitation is 300 mm, temperature range from 22,6°C to 36,3°C and an altitude from 65 to 220 meters above sea level (m.a.s.l.).

The field sampling was done during the months of February, May, and September (each sampling with a duration of five days), this period allows including rainy and wet seasons and in this way we can obtain information about the temporal and seasonal variation of the mosquito populations. In the village, we sampled two sites: one inside a patch of forest (extra domicile) and the other one in the Savannah (peri and intradomicile).

Mosquitoes were collected using active search (aspirators), CDC and Shannon traps, each method used during the same period of time (five days). The catches occurred quarterly including a diurnal period (one hour and 30 minutes before the crepuscular period), 30 minutes comprising the vespertine crepuscular period, and a following nocturnal period (one hour and 30 minutes) of mosquito activity. Resting sites such as trees, caves, rocks, and vegetation near human settlements were sampled. Sampling was developed for 5 days in each fieldwork. This period of time allows us to include the dry and wet seasons in the region.

In the field, the mosquitoes collected were killed by freezing in dry ice, sorted by genus and transported to the laboratory where they were identified according to sex and species using morphological keys (Lane 1953, Forattini 2002, Faran 1981, Chavarri 1995, Gonzalez 2009, Peccor 1992).

To characterize the populations of mosquito vectors, ecological variables were used. Thus, the following indices were calculated: (i) the ecological indices of composition: total (N), relative abundance (RA) and diversity indexes; and (ii) the ecological indices of structure: Shannon-Weaver diversity index (H′), Simpson’s diversity index, Margalef diversity index, and Sorensen equitability index. The Jaccard similarity index was used to estimate the similarity between hábitats.

## Results

A total of 1071 mosquitoes was collected and classified into the Culicidae family and two subfamilies (Culicinae and Anophelinae) and five genera: *Aedes, Culex, Psorophora, Mansonia* and *Anopheles*, and 14 species. The most diverse genera were *Culex* (5 species), *Aedes* (3 species) and *Psorophora* (2 species). The most abundant species were *Cx quinquefasciatus* (22.6%), *Ae. aegypti* (20.08%), *Ae. scapularis* (12.3%), *Ae. angustivittatus* (9%) and *An. albimanus* (8.0%); other species found in less abundance was *Phosorophora ferox* and *Mansonia indubitans*.

The traps located in the extra domicile (Shannon and CDC) were most effective (53% of the captures). The traps located in the peridomicile and intradomicile reach 20 and 27% of mosquitoes catch. The most dominant species in the extra domicile was *Ae. scapularis* followed by *Ae. angustivittatus*. The most common species in the peridomicile were *Cx. quinquefasciatus* and *An. albimanus* and the most common specie in the intradomicile were *Ae. aegypti* (Table 1).

**Table I.** Relative abundance (RA) and distribution of species in sampling sites

| Species | Forest |  | Savannah |  | Total | RA (%) |
| --- | --- | --- | --- | --- | --- | --- |
|  | Ni | pi | Ni | Pi |  |  |
| *Ae.(Och.) scapularis* | 100 | 0.107 | 44 | 0.118 | 82 | 12.03 |
| *Ae.(Och.) aegypti* | 0 | 0.000 | 240 | 0.025 | 240 | 20.08 |
| *Ae. (Och.) angustivittatus* | 70 | 0.021 | 38 | 0.054 | 22 | 3.78 |
| *An.(Nys.) albimanus* | 96 | 0.002 | 0 | 0 | 96 | 0.17 |
| *Cx. (Cux.) quinquefasciatus* | 171 | 0.084 | 100 | 0.074 | 271 | 22.6 |
| *Cx. (Cux.) nigripalpus* | 44 | 0.086 | 6 | 0.02 | 50 | 8.59 |
| *Cx.(Mel.) pedroi* | 45 | 0.088 | 0 | 0 | 45 | 7.73 |
| *Cx. (Mel.) erraticus* | 16 | 0.031 | 20 | 0.099 | 36 | 6.18 |
| *Cx.(Cx.) coronator (complex)* | 6 | 0.012 | 10 | 0.049 | 16 | 2.74 |
| *Ma. (Man) indubitans* | 1 | 0.002 | 0 | 0 | 1 | 0.17 |
| *Ma.(Man.) titillans* | 4 | 0.008 | 1 | 0.005 | 5 | 0.85 |
| *Ps.(Jan.) ferox* | 61 | 0.113 | 2 | 0.01 | 63 | 10.82 |
| **Total number of individuals (N)** | **610** | **0.808** | **461** | **0.099** | **1200** | **100** |
| **Total number of species (S)** | **11** |  | **9** |  |  |  |

In September we found the major abundance of mosquitoes, this month coincide with the highest values of temperature and relative humidity. In February (dry season) the most abundant species were *Ae. scapularis* and *Cx. quinquefasciatus*. In may (rainy season) the predominant species were *Ps*.*ferox, Cx. nigripalpus, Cx. pedroi* and *Ma. titillans*.

The estimated Margalef index for both sampling sites was 4,79 (forest) and 4,04 (savannah), by this way, the alpha diversity in both sites is similar. The Simpson index was 0,07 for the forest and 0,05 for the savannah showing a high diversity of species in the sampling and homogeneous frequency distribution.

The obtained value for the Jaccard index by comparing both sampling sites was 0,35 showing that both sites have a similar composition of species (Table 2).

The estimated Cody and Sorensen indexes for both sampling sites was of 0,22 and 0,39 respectively, indicating low replacement of species from the sites; the Shannon-Wiener index was 2,60 for the forest and 2,78 for the savannah, indicating that the diversity of species between sites is similar like was corroborated by the Jaccard index.

**Table II.**
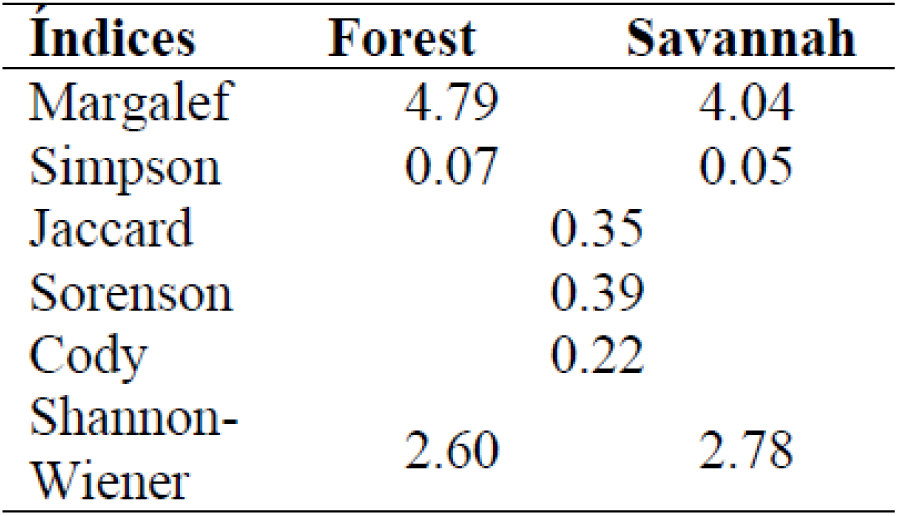
Ecological índices

The most efficient sampling methods was the active search, a method that allows to collect the 60% of the mosquitoes, followed by the CDC trap that collects the 26% of the mosquitoes. The Shannon trap was the less efficient method, 14%. With respect to the species composition by capture method, the CDC and active search methods allows to catch 9 species, the less efficient method was the Shannon trap with 4 species.

## Discussion

This is a descriptive study where we studied the dynamics of Culicidae in two different ecosystems during the same period for eight months, using ecological indices of composition and structure. To the best of our knowledge, this is the first time that such approach has been used in a settlement of indigenous communities of Magdalena State in Colombia. Also, this study is a contribution to the actualization of the inventory of species potential vectors of diseases in the zone. Of the found species, nine have been incriminated as vectors of pathogens in different regions of the Americas or Colombia. With respect to species of *Aedes* genera, they predominate in the sampling site with more conservation (forest), *Ae. scapularis* has tendency to visit the human environment and has developed some degree of sinantropy (Forattini, 2002). *Ae. angustivittatus* has been found both in sylvatic and peri and intradomiciliary environments with a tendency to bite equines. This specie has been reported naturally infected with Ilheus virus in Panamá and with the virus of Venezuelan equine encephalites in Colombia (Forattini, 2002). *Aedes aegypti* was the most predominant species in the intradomicile and is well know his capacity to transmit arbovirus such as chikungunya, dengue and zika, but the health authorities in charge of this indigenous community under-report these diseases and the community are uninformed about the risk of the presence of this mosquito specie in his environment.

With respect to the species of *Culex* genera draw attention to the finding of *Cx. quinquefasciatus* with high density in the forest because this specie is mainly found in urban environments, however, Forattini, 2002, was reported his presence in the rural environment, this finding demonstrates that is an eclectic species not only restricted to the human environment.

*Cx. nigripalpus* is a very versatile species for choosing habitats. In the present study predominated in the forest and is know that his antropophily is low and is mainly an ornithopilic specie, but has the potential to adapt to the anthropic environment (Forattini, 2002, Guimaraes, 2000). *Cx. nigripalpus* was incriminated as vector of San Luis Encephalites in United States (Day, 1999) and has been found naturally infected with SLE virus in Central America, Trinidad and Tobago and Ecuador (Nayar, 1968; Tsai, 1989).

For West Nile Virus has been proved the dispersion of this disease through the Caribbean region and the main vectors are *Cx. pipiens, Culex restuans* and *Cx. quinquefasciatus* (Koné, 2003). *Cx. quinquefasciatus* is of epidemiologic importance due his incrimination as vector of filariasis and arboviruses.

In Colombia recent studies (Mattar, 2005) in some departments of the Caribbean region (Córdoba and Sucre) demonstrated seroprevalence for this virus in equines. *Cx. quinquefasciatus* is considered a potential vector of WNV in the studied zones. In the research area, due to agricultural expansion and wood exploitation, wide areas of natural vegetation has been replaced for deforested zones. The mosquitoes are sensitive to the environmental changes presenting adaptation responses, these responses are reflected in the composition and abundance of the species (Gomes, 2007, Parra-Henao 2012). The diversity and abundance of mosquitoes vectors of diseases in the zone are high as shown by the alpha and beta diversity indexes. The ecological analysis plus the previous reports of vectorial capacity for some species registered in this study allows concluding that in the zone could be present the interaction between the elements of the transmission chain and thus outbreaks of vector-borne diseases could be expected and is recommended to increase the entomological vigilance. This study contributes to existing knowledge regarding the relationship between vector dynamics and drivers in Colombia. This information is critical when planning surveillance and prevention activities in zones where resources are limited.

## Acknowledgements

To Departamento Administrativo Nacional de Ciencia y Tecnología de Colombia ‘‘Francisco José de Caldas– COLCIENCIAS, Colombia for the financial support, contract number 767-2016, project code 1415744560123256-04-18067.

